# Mesenchymal-epithelial transition in lymph node metastases of oral squamous cell carcinoma is accompanied by ZEB1 expression

**DOI:** 10.1101/2022.02.03.478962

**Authors:** Kai Horny, Christoph Sproll, Lukas Peiffer, Frauke Furtmann, Patricia Gerhardt, Jan Gravemeyer, Nikolas H. Stoecklein, Ivelina Spassova, Jürgen C. Becker

**Affiliations:** Translational Skin Cancer Research, German Cancer Consortium (DKTK), 45141 Essen, Germany; German Cancer Research Center (DKFZ), 69120 Heidelberg, Germany; Department of Oral- and Maxillofacial Surgery, Medical Faculty, University Hospital of the Heinrich-Heine-University Düsseldorf, Germany; Department of Dermatology, University Medicine Essen, 45141 Essen, Germany; Department of General, Visceral and Pediatric Surgery, Medical Faculty, University Hospital of the Heinrich-Heine-University Düsseldorf, Düsseldorf, Germany

**Keywords:** single cell RNA, oral cavity, squamous cell carcinoma, Epithelial-mesenchymal plasticity, EMT, MET, ZEB1, heterogeneity

## Abstract

**Background:** Oral squamous cell carcinoma (OSCC), an HPV-negative head and neck cancer, frequently metastasizes to the regional lymph nodes but only occasionally beyond. Initial phases of metastasis are associated with an epithelial-mesenchymal transition (EMT), the consolidation phase is associated with mesenchymal-epithelial transition (MET). This dynamic is referred to as epithelial-mesenchymal plasticity (EMP). While it is known that EMP is essential for cancer cell invasion and metastatic spread, less is known about the heterogeneity of EMP states within a tumor and even less about the heterogeneity between the primary and metastatic lesions.

**Methods:** To capture heterogeneity of EMP states in OSCC, we performed single-cell RNA sequencing (scRNAseq) of 5 primary tumors and 9 matching lymph node metastases and re-analyzed publicly available scRNAseq data of 9 additional primary tumors. To account for possible bias in cell type compositions by scRNAseq, these were also deconvoluted from bulk transcriptome analyses. Protein expression of selected genes were confirmed by immunohistochemistry.

**Results:** From the 23 OSCC lesions the single cell transcriptome of a total of 7,263 carcinoma cells was available for in-depth analyses. We initially focused on one lesion to avoid inter-patient heterogeneity as a confounding factor and identified OSCC cells expressing genes characteristic of different epithelial and partial EMT stages, such as keratins and *SPRR1B* (cornifin B) or vimentin and matrix metallopeptidases. RNA velocity information together with the increase in inferred copy number variations indicated a progressive trajectory towards epithelial differentiation in this metastatic lesion. Extension to all samples revealed a less stringent but essentially similar pattern. Interestingly, cells undergoing MET show increased activity of the EMT activator ZEB1. Immunohistochemistry confirmed that ZEB1 was co-expressed with the epithelial marker cornifin B in individual tumor cells - more frequently in lymph node metastases. The lack of E-cadherin mRNA expression suggests this is a partial MET.

**Conclusions:** This study reveals that EMP enables different partial EMT and epithelial phenotypes of OSCC cells, which are endowed with capabilities essential for the different stages of the metastatic process, including maintenance of cellular integrity. During MET, ZEB1 appears to be functionally active, indicating a more complex role of ZEB1 than mere induction of EMT.

## Background

Head and neck squamous cell carcinoma (HNSCC) is the sixth most common cancer worldwide, with 890,000 new cases and 450,000 deaths in 2018 (1). The survival for HNSCC patients has improved modestly over the past decades; however, this improvement is partially attributable to the emergence of human papillomavirus (HPV)-associated HNSCC which has a better prognosis than HPV-negative tumors. One of the HPV-negative HNSCC subtypes is oral cavity squamous cell carcinoma (OSCC) which is mainly associated with tobacco and alcohol abuse (1). OSCCs are often diagnosed at an early stage owing to the patient’s self-identification of the mass lesion and symptoms, still regional lymph node metastases are frequent; thus, surgical removal of primary tumor is accompanied by neck dissection and radiotherapy (2). Given the morbidity associated with this combined intervention, there is a need to identify molecular biomarkers to predict the presence of lymph node metastases and to prognosticate survival.

Invasion and metastasis becomes possible in many epithelial tumors through an epithelial-mesenchymal transition (EMT), i.e., a reactivation of an embryonic developmental program in which cells acquire migratory and invasive properties (3). In EMT, epigenetic, transcriptional, and post-translational changes cause epithelial cells to break down the strong homotypic cell-cell junctions and adopt a mesenchymal morphology (4). It is important to note that EMT should not be understood as a clearly defined process, but rather as many dynamic and complex processes, which may vary depending on tumor entity, stage, and microenvironment (4-6). Thus, expression of EMT-related genes and their regulating transcription factors is highly heterogeneous even within one cancer entity, between patients, different lesions from one patient and between individual cancer cells within one lesion (4, 7). Since it is a continuous, dynamic, and reversible process cancer cells can adopt a multitude of intermediate or partial states, e.g., epithelial to more mesenchymal or partial EMT (pEMT) states (5, 8-11). Therefore, it has recently been recommended that this EMT continuum should rather referred to as epithelial-mesenchymal plasticity (EMP) (4, 10).

To capture the EMP-associated heterogeneity of cancers, single-cell analyses appear to be the most appropriate approach, however, to date most EMP-related single-cell studies are based on controlled *in vitro* and *in vivo* experiments (5, 6, 9, 12, 13). Particularly in HNSCC only few studies scrutinized EMP within freshly isolated tumor samples (8, 14, 15). Especially noteworthy is the landmark paper by Puram *et al*. They characterized 2,215 malignant cells from 18 patients and discovered several pEMT states with high variability in EMP-related gene expression (8).

In the work presented here, we investigated the phenotypic heterogeneity of 5 primary and 9 regionally metastatic OSCC lesions isolated from 7 patients using multiplexed single-cell RNA sequencing (scRNAseq). In addition, we used a recently published series of scRNAseq data from primary HNSCC that included 9 OSCC tumors to put our observations on an even broader data base (15). Our results not only confirm the EMP-associated heterogeneity of cancer cells in primary and metastatic OSCC, but also demonstrate mesenchymal-epithelial transition (MET) in established lymph node metastases. Surprisingly, we observed a high activity of EMT-activator ZEB1 in metastatic OSCC cells with epithelial differentiation, a notion which was confirmed by co-expression of ZEB1 and cornifin-B protein in individual tumor cells.

## Methods

### Tissue samples

19 tissue samples that included 5 primary tumors, 14 potentially affected lymph node metastases, from which 9 represent lymph node metastases as we detected carcinoma cells in histopathological examination and scRNAseq from 7 OSCC patients treated at the Department of Oral-, Maxillo- and Plastic Facial Surgery, University Hospital of the Heinrich-Heine-University Düsseldorf, Germany, were used for histology, IHC, immune fluorescence as well as bulk and single-cell transcriptome analyses. The clinical details are provided in table 1.

### Histology and IHC

Hematoxylin and Eosin (H&E) and IHC were performed on 4 μm formalin-fixed, paraffin-embedded (FFPE) sections. H&E staining was performed using standard protocols (Supplementary Figure 1). Whole-slide imaging was performed using Zeiss Axioscan 7 and 10x magnification (Carl Zeiss Microscopy Deutschland GmbH, Oberkochen, Germany).

IHC using the rabbit polyclonal antibodies anti-SPRR1B (Cat. No.: SAB1301567-400UL, Sigma Aldrich, Darmstadt, Germany) anti-ZEB1 antibody (Cat. No. HPA027524-25UL, Sigma Aldrich) was performed as previously described (16). Briefly, after sections were deparaffinized for 60 min at 60°C and rehydrated, sections were incubated for 15 min in an inverter microwave oven with antigen retrieval buffer pH 9 for anti-SPRR1B and pH 6 for anti-ZEB1. After 3×2 min washes with Tris-buffered saline with 0.1% Tween (TBST) sections were incubated for 8 min with 3% peroxidase and after an additional washing step for 30 min with 3% bovine serum albumin (BSA) in TBST. For single staining sections were subjected for 1h to anti-SPRR1B in 1:600 dilution or overnight to anti-ZEB1 in a 1:500 dilution, both at room temperature. Afterwards, the secondary anti-rabbit HRP Polymer was applied for 30 min, followed by 1:20 diluted 3,3’-Diaminobenzidin (DAB) for 10 min and 1:10 diluted hematoxylin for 3 min with washing steps with TBST in between before a final washing with tap water for 3 min before fixation. For multiplexed antigen detection, the OpalTM chemistry system (Akoya Biosciences, Marlborough, MA, USA, Cat. No.: OP7TL4001KT) was used according to the manufacture’s description. Briefly, after deparaffinization and fixation, we processed the sections for 15 min with retrieval buffers in an inverter microwave oven. Then, we incubated them with antibody diluent for 10 min at room temperature, followed by incubation with the anti-SPRR1B antibody for 30 min. Next, Opal Polymer horseradish peroxidase (HRP) secondary antibody solution with the respective chromogen was applied for 10 min, antibodies were removed by microwave treatment and the staining with anti-ZEB1 antibody was performed. Slides were lastly incubated with 4′,6-diamidino-2-phenylindole (DAPI) for 5 min.

### Single-cell RNA sequencing

Samples were processed immediately after surgery and temporarily stored for transport at 4°C in tissue storage solution (Miltenyi Biotec, Bergisch Gladbach, Germany) as previously described. Briefly, samples were dissociated into single-cell suspensions using the gentleMACS Dissociator (Cat. No. 130-093-235, Miltenyi Biotec, Bergisch Gladbach, Germany) with program “h_tumor_01”, followed by 2x program “h_tumor_02” in 4.7 ml RPMI 1640 (Cat. No. P04-16500, PAN-Biotech) and an enzyme mix consisting of 200μl Enzyme H, 100 μl Enzyme R and 25 μl Enzyme A (Cat. No. 130-095-929 Miltenyi Biotec). Afterwards, single-cell suspensions were reconstituted and washed thrice with 0.05 % BSA phosphate-buffered saline (PBS) and filtered through a 100 μl cell strainer.

In cases multiple samples of one patient had to be analyzed (Table 1), antibody hashing for multiplexing of samples was performed according to manufacturer’s protocol. Briefly, 1 μg of the respective TotalSeq anti-human hashtag antibody was used to incubate a maximum of ca. 2 million cells for 30 min at 4 °C (Cat. No. 394601, 394603, 394605 and 394661, 394663, 394665, respectively, Biolegend, San Diego, CA, USA). After 3 washes with PBS with 0.05 % BSA respective cell suspensions were mixed prior to single-cell RNA library preparation. In short, both unhashed and hashed single-cell suspensions were barcoded and processed with the microfluidic system of 10x Genomics Chromium v2.0 platform as described in the manufacturer’s protocols (10x Genomics, Leiden, Netherlands). Due to a change of system, both the 3’ technology including Chromium Single Cell 3’ Library & Gel Bead Kit version 2 (Cat. No. 120237), Chromium Single Cell A Chip Kit (Cat. No. 120236), Chromium i7 Multiplex Kit (Cat. No. 120262) as well as the 5’ technology including Chromium Single Cell 5’ Library & Gel Bead Kits version 2 (Cat. No. 1000263), Chromium Next GEM Chip K Single Cell Kit (Cat. No. 1000286), Dual Index Kit TT set A (Cat. No. 1000215) were used; for library construction the Chromium Single Cell 3’/5’ Library Construction Kit (Cat. No. 1000020) was applied. After library preparation, the library from patient one was sequenced with an Illumina HiSeq 4000 (Illumina, Berlin, Germany) at the DKFZ Genomics and Proteomics Core Facility in Heidelberg and all other libraries were sequenced on an Illumina NovaSeq 6000 (Illumina, Berlin, Germany) in three runs (Run 1: patient 2,4,5; Run2: patient 3, Run3: Patient 6 and 7) at the West German Genome Center in Cologne.

### Bulk transcriptome analysis

The bulk transcriptome was analyzed using a quantitative nuclease protection assay from the HTG Transcriptome Panel (HTP) according to the manufacturers protocol (Cat. No. HTG-001-224, HTG Molecular Diagnostics, Tucson, AZ, USA). Briefly, the tumor areas were macro-dissected as depicted in Supplementary Figure 1 from 4 μm FFPE sections and subjected to Proteinase K and DNase digestion. Next, the quantitative nuclease protection assay was performed using the HTG EdgeSeq Processors before adapters and sample tags were added in a PCR amplification. The resulting libraries were sequenced using on an Illumina NextSeq 500/550 High Output Kit v2.5 (75 cycles) (Cat. No. 20024906, Illumina, Berlin, Germany).

The resulting FASTQ files were processed towards a gene expression count matrix using the HTG EdgeSeq Reveal Software version 4.0.1. Quality Control, normalization, and principal component analysis (PCA) were performed using R version 4.0.5. Sample 5 failed QC due to low number of sequenced reads and was removed from the analysis. Deconvolution was performed with web application of CIBERSORTx (https://cibersortx.stanford.edu/) using a signature matrix derived from the gene expression count matrix of combined scRNAseq data of the samples analyzed with HTP (17). We filtered for genes that are expressed less than 5% within the given tumor phenotypes and randomly included only 75% of T- and B-Cells for better performance. The resulting signature matrix was used for imputing cellular fractions from the counts-per-million normalized HTP data without any batch correction or quantile normalization and 500 permutations.

### Bioinformatic analysis of scRNAseq data

#### Preprocessing

Processing from FASTQ files towards the unfiltered count matrix (barcodes x genes) was performed using Cellranger Software Suite version 3.1.0 and the human reference genome build GRCh38, downloaded from 10x Genomics in version 3.0.0.

Cells were identified by evaluating quality criteria inspired by Luecken *et al*., also see Additional file 1 (18). Cells were defined by having more than 500 UMIs, less than 10 % mitochondrial gene expression and additionally for patient 3 and 5 having more than 30 housekeeper genes expressed. The filtered count matrices (cells x genes) were further processed using Seurat version 4.0.1 and R version 4.0.5 (19). Demultiplexing of hashed libraries was performed choosing manual threshold of hashtag oligo (HTO) expression based on quality assessments described in Additional file 1. We removed doublets that were identified by demultiplexing of HTO expression matrices from all analyses.

#### Normalization and dimensionality reduction

Every time we performed analysis of a specific set of cells, e.g., only tumor cells or only a specific patient, the respective set of cells is normalized using the SCtransform algorithm and the 3000 most variable genes were selected for PCA (20). During normalization, we regressed for cell cycle scores and percentage of mitochondrial gene expression. Cell cycle scores and phases were determined in Seurat using log-normalized RNA counts and S- and G2M-Phase genes defined by Tirosh *et al*. (21). When generating a uniform manifold approximation and projection (UMAP) without patient-specific batch effect we used corrected PCs derived with the harmony R package version 0.1.0 [54]. Based on the variance explained by each PC and the respective ranked elbow plot we choose an appropriate number of PCs for UMAP visualization and SNN clustering as implemented in Seurat. For deriving patient-specific clusters for calculating the intratumoral cosine similarity we used the same resolution parameter for comparability. UMAPs colored by specific gene expression were ordered by expression values.

#### Tumor cell identification and phenotyping

We annotated all cell types and identified tumor phenotypes by using a combination of methods: SNN clustering, differential gene expression, gene set enrichment analysis (GSEA), expression of literature-based marker genes and automated reference-based annotation with SingleR using the Monaco bulk-RNA Immune dataset (22, 23). Automated, reference-based annotation with SingleR version 1.4.1 was run on SNN clusters with resolution 100 yielding extremely small cluster including only few cells but increasing performance. Further, we excluded cells with ambiguous cell type marker expression. For example, within the tumor cells of the lymph node metastasis from patient 1, we observed few cells expressing genes typical for fibroblasts and DCs that were subsequently excluded from tumor-specific analysis. Similarly, we removed T-Cells, B-Cells, mast cells, fibroblasts, muscle cells, melanocytes, and other Immune cells from the tumor cell subsets. Malignant cells were first identified by high epithelial gene expression as for example cytokeratins and inferred CNVs (Supplementary Figure 2). CNVs were inferred using the R package inferCNV version 1.6.0 with the not normalized, filtered count matrix including all cells as input and algorithm run in “samples” mode. Inferred CNVs of mitochondrial genes were excluded. Differential gene expression was performed by calculating the log2 foldchanges between one cluster and all other cells from the subset based on log-normalized data using NormalizeData function and scale factor of 10,000. We filtered for genes with log2 foldchange greater than 0.25 and a minimum percentual expression of at least 10 % within the cluster or all other cells. For calculating the cosine similarity, we did not filter the log2 foldchanges. GSEA was performed using the “fgsea” R package version 1.16.0, log2 foldchanges from differential gene expression and gene ontology biological processes (GO:BP,C5 v7.1) as well as hallmark gene sets (H, v7.1) downloaded from MSigDB database (24-26). Gene sets were included if they have at least 15 genes or at maximum 500 genes within the gene set using 10,000 permutations.

For deriving epithelial differentiating and pEMT gene signatures of patient 1 we calculate foldchanges between the “epi” and “pEMT” cluster and included genes with log2 foldchanges greater or lesser than 1, respectively, and with at least 10 % of either epi or pEMT cells expressing that gene. Gene set variation analysis (GSVA) was performed using the R package GSVA version 1.38.2 with default settings, i.e., gaussian kernel (27). As input, EMTome signatures, the pEMT and epithelial differentiation 1 and 2 signature from Puram *et al*. (8) and the three EMT and the epithelial senescence signatures from Kinker *et al*. were used (12).

Trajectories were inferred using SlingShot version 1.8.0 with malignant cell clusters as shown in Figure 2A of patient 1 and the first 20 PCs (28). RNA velocity was inferred using the VeloCyto python and R package (version 0.6) (29). Creation of the loom file was done using default options and gene annotations as used for Cellranger processing. Genes were filtered based on the minimum average expression magnitude with a threshold of 0.05 for spliced and 0.02 for unspliced reads. Velocity estimates were calculated using as distance the inverse correlation coefficient of the PC embedding correlation matrix, the top/bottom 2 % quantiles for gamma fit, 50 neighboring cells projecting 1 deltaT into the future and projected on the UMAP using 200 neighbors and 50 grid points.

Transcription factor activity was inferred using the VIPER algorithm (version 1.24.0) and regulons from the DoRothEA database (version 1.2.2) (30-32). Hierarchical clustering in the heatmaps was performed using Euclidean distance and ward.D2 method unless otherwise noted. Visualization was performed using ggplot2 version 3.3.3 and ComplexHeatmap version 2.6.2 (33, 34).

### Reanalysis of HNSCC dataset from Kürten *et al*

From the publicly available scRNAseq data set on primary HNSCC tumors we downloaded the FASTQ files of CD45-negative and HPV-negative libraries from the sequencing read archive (SRA) under accession ID SRP301444 (15). The data sets were analyzed the same as described above. HPV-negative samples were chosen for comparability to the OSCC dataset. However, the HN07 tumor originated from the larynx, while all other samples indeed originated from the oral cavity. We adjusted the cell identification thresholds based on our evaluation criteria pooled for all libraries: cells were defined by having more than 175 genes expressed, less than 10 % mitochondrial gene expression and more than 60 housekeeper genes expressed (see Additional file 1).

## Results

### Single-cell gene expression signatures of tumor cells from a single metastasis show several predominant, but not necessarily exclusive, functional phenotypes

To avoid inter-patient heterogeneity as a confounding factor, we first focused on the analysis of a single OSCC metastasis to develop hypotheses which would be subsequently tested in the entire cohort. For this, we chose a metachronous lymph node metastasis that was removed one year after the primary tumor, because we assumed that consolidation processes are particularly pronounced in this longer existing metastasis. Multiplexed scRNAseq recovered 4,121 cells that could be assigned to the following cell types: 1,906 (46.8 %) tumor cells, 1,186 (29.1 %) fibroblasts, 507 (12.4 %) dendritic cells (DCs), 375 (9.2 %) macrophages and 102 (2.5 %) endothelial cells (ECs) (Figure 1A). The absence of T- or B-cells was in line with the histology showing completely disrupted lymph node structures and only occasional tumor-infiltrating lymphocytes (Supplementary Figure 1). OSCC cells were identified both by the presence of copy number variations (CNVs) inferred from scRNAseq data as well as the expression of epithelial markers including *S100A2, S100A14*, cytokeratins (*KRT5, KRT14, KRT17)* and stratifin (*SFN*) (Supplementary Figure 2).

**Figure 1.**
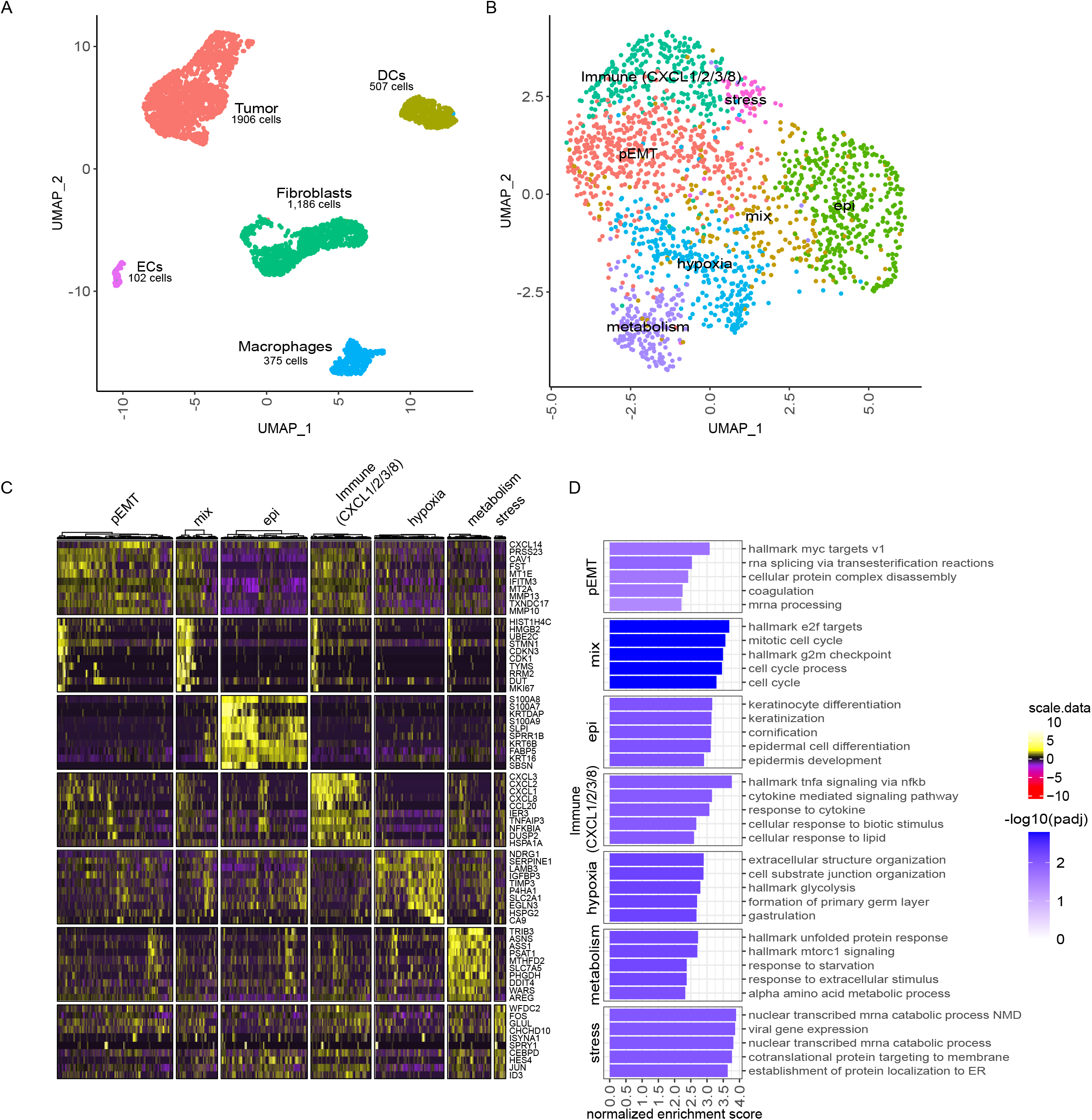
Single-cell gene expression signatures in OSCC cells from a single metastasis reveals predominant functional phenotypes. **(A)** UMAP based on scRNAseq data of 4,076 cells isolated from a metachronous lymph node metastasis. Cells are annotated and summarized according to the presumed cell type. **(B)** UMAP of 1,906 OSCC cells depicted in A. Cells are annotated according to predominant functional phenotype. **(C)** Heatmap for scaled, log-normalized gene expression of tumor cells (columns) split by respective phenotype and the top 10 differentially expressed genes (DEGs) (rows) of the respective phenotype against all other tumor cells. DEGs are sorted from highest to lowest log2 foldchange. Row sections are ordered like column sections. **(D)** Top 5 enriched gene sets from log2 foldchanges of respective tumor phenotypes by normalized enrichment scores (x axis). Gene sets of respective phenotypes are sorted from highest to lowest enrichment. Bars are colored by the negative decadic logarithm of the Benjamini-Hochberg adjusted p-value (padj). DCs: dendritic cells. ECs: endothelial cells.

Detailed phenotyping of the cancer cells identified several clusters to which we could assign the following predominant functional phenotypes that differ in their EMT state, immunomodulatory capacity, as well as their response to hypoxia, stress, and metabolic constraints (Figure 1B-D). However, the predominance of a functional phenotype does not preclude additional traits. Specifically, 515 cells exhibited a pEMT phenotype by expressing a mixture of epithelial and mesenchymal genes such as matrix metallopeptidases (*MMPs*), caveolin-1 (*CAV1*) and *CXCL14* that were previously described in pEMT signatures and are enriched in the EMT hallmark gene set from the molecular signatures database (MSigDB) (Supplementary Figure 3A) (8, 25). In contrast, 385 cells showed higher expression of genes associated with epithelial differentiation such as *S100A7/A8/A9*, the keratinocyte envelope protein cornifin-B (*SPRR1B*), and various cytokeratins (e.g., *KRT6B* and *KRT16*). The EMP-related gene expression patterns are correlating with established signatures from the EMTome database (Supplementary Figure 3B-E) (7). Interestingly, both gene expression signatures were present in a mixed cluster with 184 cells characterized by the high expression of cell-cycle related genes (despite the fact that we applied cell cycle regression).

With respect to cell clusters whose gene expression was not predominately associated with EMP, 268 cells exhibited an immune-regulatory phenotype enriched for genes associated with cytokine-mediated responses and higher expression of the chemokines *CXCL1/2/3/8* and *CCL20* and for 554 cells the gene expression pattern suggests adaptation to environmental factors. In detail, 51 cells had higher expression of transcription factors *FOS* and *JUN* suggesting a stress response. 310 cells can be assumed to respond to hypoxic conditions in the tumor based on the higher expression of *NDRG1* and *EGLN3* both of which are regulated by oxygen levels (35, 36): *NDRG1* regulates stress response and p53-mediate caspase activation (35), and *EGLN3* has an important role in hypoxia-inducible factor 1 alpha (HIF1α) regulation through prolyl hydroxylation (36). 193 cells expressed genes associated with amino acid metabolism, starvation response and mTORC1 signaling, i.e., a regulator of mitochondrial metabolism (37). The upregulated genes *ASNS, PSAT1* and *PHGDH* integrate the metabolites of serine and glycine metabolism into glycolysis and therefore fuel glycolysis with amino acids (38); hence, these cells appear to be adapted to low-glucose conditions.

### OSCC cells in lymph node metastases undergo mesenchymal-epithelial transition

We next focused on possible dynamics within the predominantly EMP-related cancer cell clusters using a higher resolved shared-nearest neighbor (SNN) clustering. We defined 4 pEMT clusters (pEMT-1 to 4), 4 clusters of more epithelial differentiated cells (epi-1 to 4) and one cluster with mixed phenotypes (mix) (Figure 2A). pEMT-1 is enriched for genes involved in coagulation and play a role in angiogenesis as *THBS1, CYR61* and *F3* (Figure 2B, Supplementary Figure 4A). pEMT-2 and 3 showed higher expression of extracellular matrix (ECM) remodeling genes, but pEMT-2 also higher expressed cytokeratin *KRT15* and chemokine *CXCL14* while pEMT-3 higher expressed the serine protease inhibitor *SERPINA1* and podoplanin (*PDPN*) which mediates efficient ECM degradation by controlling invadopodia (39). Of the more epithelial differentiated cell clusters, epi-2’s expression profile is closest to the pEMT cluster: higher expression of *MMP1* and lower expression of *SPRR1B* and *S100A8/A9* than epi-1, 3 and 4 (Figure 2B, E). Epi-3 showed higher expression of *S100A7* and *KRTDAP*, whereas epi-4 has higher expression of kallikreins (*KLK6/7*), prostate stem cell antigen (*PSCA*) and adipogenesis regulatory factor (*ADIRF*). Both *ADIRF* and *PSCA* play a role in prostate cancer and *PSCA* is also reported as highly expressed in mucosal tissue, but less in HNSCC (40-42).

**Figure 2.**
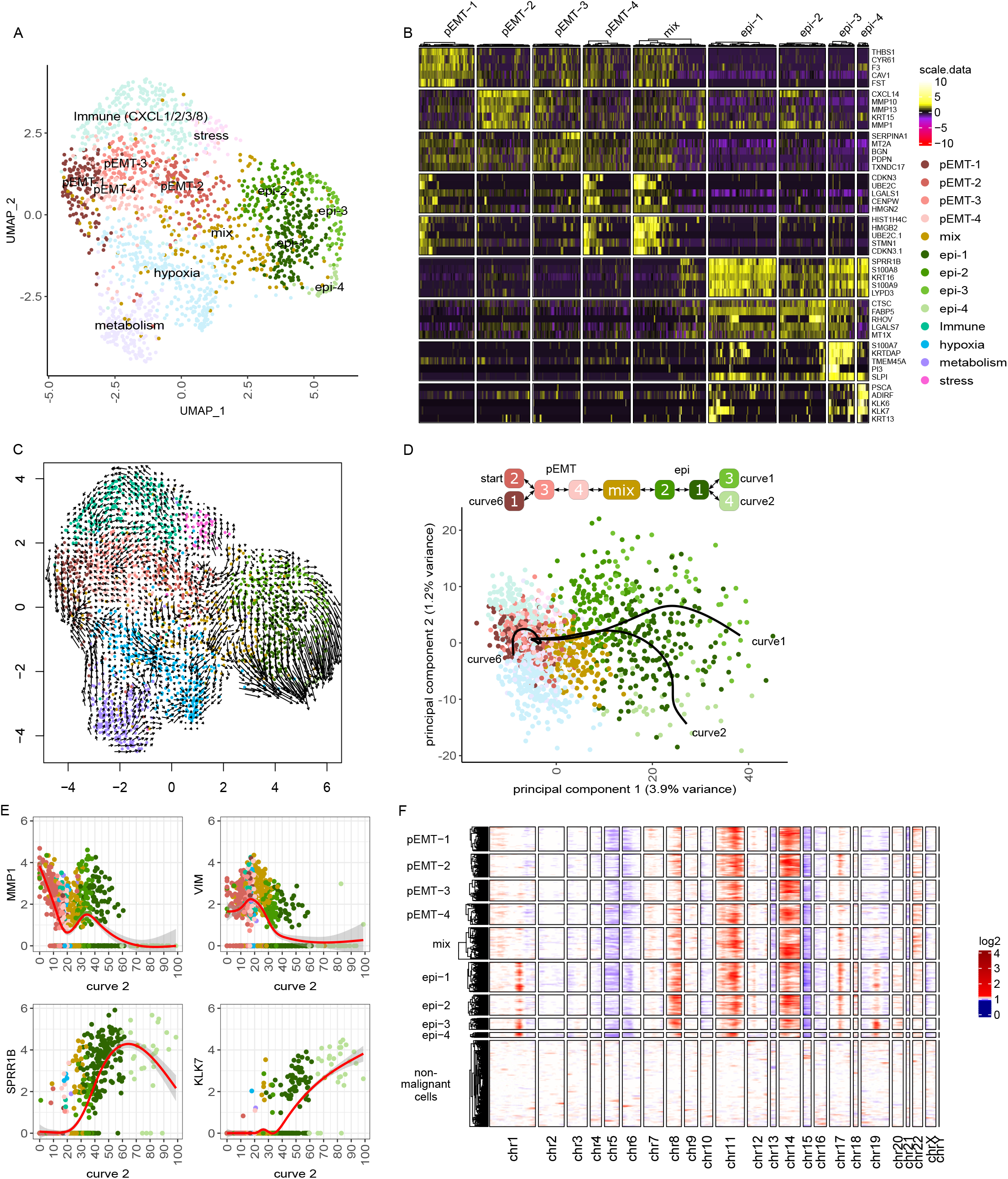
Progressive epithelial differentiation, but no strong uniform direction of development in pEMT clusters. **(A)** UMAP of 1906 OSCC cells annotated based on SNN clustering, defining 4 pEMT (pEMT-1 to 4), 4 epithelial differentiated (epi-1 to 4) and one mixed (mix) cluster; clusters are numbered by size. **(B)** Heatmap for scaled, log-normalized gene expression in EMP-associated tumor cell phenotypes (columns) split by EMP cluster and their top 5 DEGs (rows) against all other EMP-related tumor cell phenotypes. DEGs are sorted from highest to lowest log2 foldchange. Row sections are ordered like column sections. **(C)** Projection of RNA velocity on the UMAP depicted in A. Arrows indicate the extrapolated direction of development and their length the strength of this future development. **(D)** First two principal components of OSCC cells with the three EMP-related principal curves that are derived from trajectory inference. Graph on top visualizes the relationship between EMP clusters described by the three principal curves forming a branching trajectory. **(E)** Log-normalized expression (y-axis) of MMP1, VIM, SPRR1B and KLK7 across pseudotime values (x-axis) of curve 2, color-coded by clusters. Red lines indicate smoothed expression values over the trajectory generated with a general additive model; 95 % confidence intervals are shaded gray. **(F)** Inferred CNVs across EMP-related tumor cells (rows) for all chromosomes (columns). Red indicates copy number gains, white diploid copy number and blue copy number loss. Columns show genes categorized in chromosomes and ordered by genome position; hence the size of the chromosome reflects the number of detected genes and not its nucleotide length. Mitochondrial genes were excluded.

To gain a better understanding on the gene expression dynamics we estimated RNA velocity, which predicts the short-term future development in gene expression of individual cells using the ratio of spliced and non-spliced mRNA counts (Figure 2C). This analysis revealed that epithelial differentiated cells were strongly developing towards cluster epi-4, while most other cells did not show a strong uniform developmental direction. Tracking the developmental pathway within the metastasis by trajectory analysis across all EMP-related clusters revealed a major developmental axis between pEMT and epithelial differentiated cells that is diversifying within each end (Figure 2D, E, Supplementary Figure 4B). To confirm that we the found progressive epithelial differentiation of the metastatic cells represents a MET, we inferred CNVs from the scRNAseq data. This demonstrated an increased number of copy number gains on chromosome 1, 8, 17, and 19 within epithelial differentiated cell clusters epi-1, epi-3, and epi-4 (Figure 2F, Supplementary Figure 4C). It should be noted, however, that CNVs on chromosomes 1 and 17 are associated with upregulated epithelial genes in close genomic proximity; thus, these two copy number gains may not represent true genomic CNVs, but rather reflect the high expression of these genes in epithelial differentiated cells (Supplementary Figure 4D, E).

### Intra-tumoral heterogeneity of OSCC is driven by EMP

To test whether our observation that OSCC cells in one lymph node metastasis undergo a mesenchymal-epithelial transition is generally valid we extended our analyses by adding 5 primary tumors and 8 matched lymph node metastases from 6 patients (Table 1). From the publicly available scRNAseq data set on primary HNSCC tumors published by Kürten *et al*. (15) we chose to include the 9 HPV-negative primary tumors, which were all but one originating from the oral cavity (HN07 originated from the larynx) in our analysis. In total, we analyzed 7,263 cancer cells from 16 different patients (Figure 3A). Importantly, the frequency of cancer cells was unevenly distributed across samples, which could not be explained by differences in tumor cell content across samples as determined by histopathology (Supplementary Figure 1). Since the recovery of cells by scRNAseq is also low, the stability of epithelial tumor cell assemblies, which may not be sufficiently broken up by dissociation protocols, likely interfered with the generation of OSCC single-cell suspensions. In addition, as expected from the inter-patient heterogeneity, cancer cells were clustered based on their gene expression by patient rather than functional phenotype (Figure 3A, B). Thus, we accounted for the patient-specific effects with batch-corrected principal components (PCs) using the harmony R package which indeed resulted in a clustering by functional phenotypes (43) (Figure 3C, Supplementary Figure 5A-C). For initial annotation of the phenotype of each cluster, we were guided by the gene signatures previously identified in the indicator sample, with additional predominantly immunoregulatory phenotypes found. EMP-related phenotypes were present in all, but one tumor sample with only one cell (Figure 3D). Nevertheless, direct comparison of the frequencies of the predominant phenotypes in the pooled scRNAseq dataset proved difficult to interpret due to differences in cell numbers, and patient-specific gene expression patterns. To confirm the validity of our scRNAseq data, we examined the bulk transcriptome and deconvoluted the respective cell types for our 18 OSCC samples (Supplementary Figure 6). Using this approach, we observed a higher tumor and stroma content and less lymphocytes, which was expected as scRNAseq is known to be biased towards leukocytes. Therefore, we chose an alternative approach: to compare the EMP-related intrapatient heterogeneity of all analyzed tumor cells between patients, we calculated the similarities of the differential gene expression within each patient of all respective clusters of all patients. We first considered each patient individually and performed clustering, annotation, inferCNV and differential expression analysis. As exemplified for the primary OSCC of patient HN01, tumor cell clusters had the same inferred CNVs and were annotated based on their phenotype using again the indicator sample as a guide (Figure 3E, Supplementary Figure 5D, E). The cosine similarity between patient-specific clusters demonstrated that within each patient, the heterogeneity in EMP is most prominent and epithelial differentiated phenotypes are profoundly different from most other, especially pEMT clusters (Figure 3F, Supplementary Figure 5F). Indeed, all pEMT phenotypes are very similar to each other and show a large overlap of the gene expression patterns with predominantly immune- and metabolic-related clusters. In epithelial differentiated cells, the most upregulated genes are *S100A8* and *S100A9*, encoding calprotectin, and *SPRR1B*, all members of the epidermal differentiation complex (44). Of note, hypoxia- and stress-related heterogeneity is similar between patients suggesting a reactive response rather than an aspect of tumor evolution.

**Figure 3.**
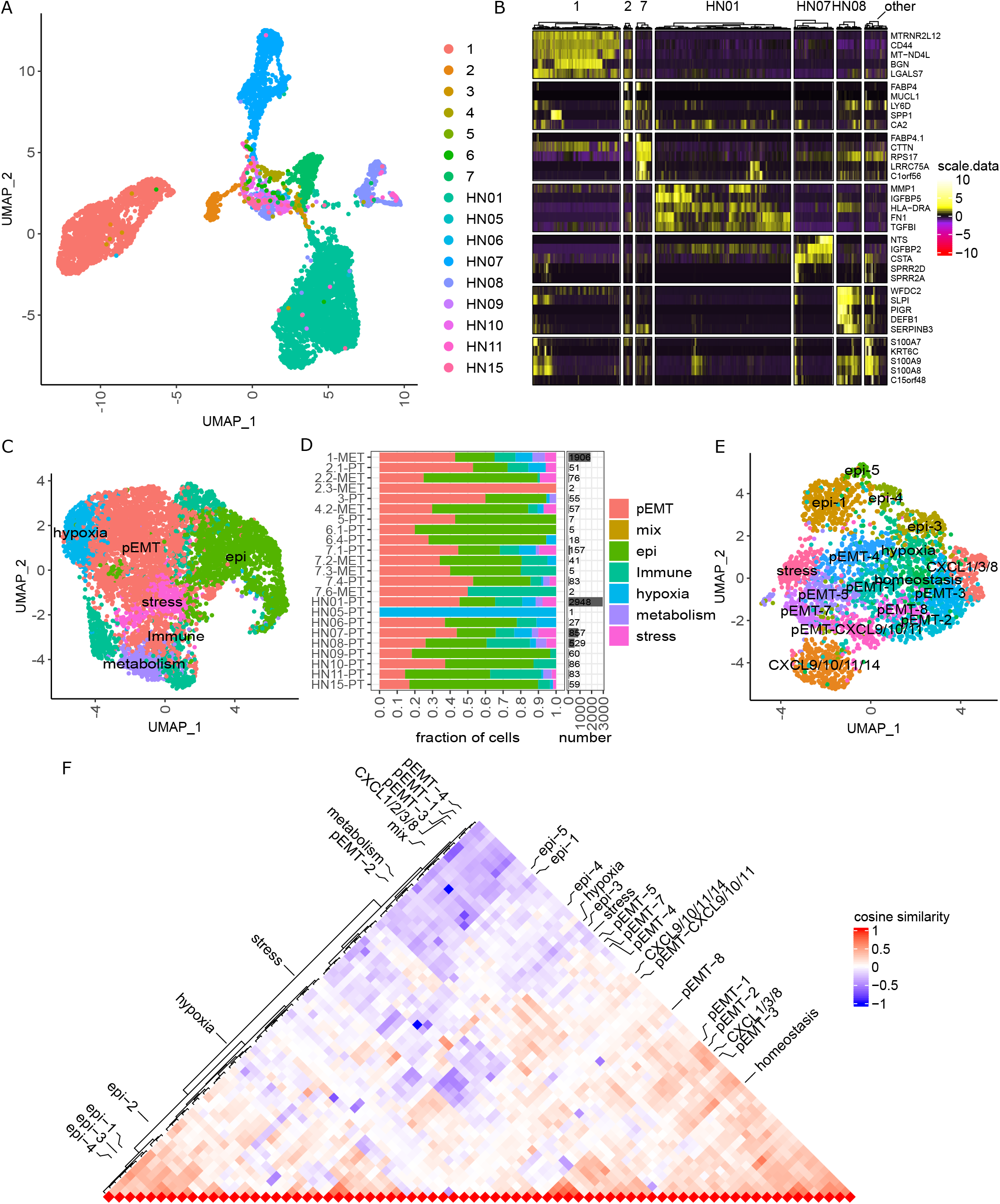
Intra-tumoral heterogeneity of OSCC is driven by EMP. **(A)** UMAP based on scRNAseq data of 7,263 cancer cells from 16 different patients annotated by patient. **(B)** Heatmap for scaled, log-normalized gene expression of tumor cells split by patients and their top 5 DEGs (rows) against all other tumor cells. All patients with less than 100 cells are summarized in the ‘other’ column. DEGs are sorted from highest to lowest log2 foldchange. Row sections are ordered like column sections. **(C)** UMAP based on scRNAseq data depicted in A with PCs corrected for patient-specific effects using harmony. Cells are annotated according to their predominant phenotype. **(D)** Relative distribution of tumor cell phenotypes (left) and cancer cell abundance (right) across patients. The label on the y-axis shows the sample identification and tumor localization (primary tumor [PT] or lymph node metastasis [MET]). **(E)** UMAP based on scRNAseq data of 2,948 OSCC cells from patient HN01. Cells were annotated based on SNN clustering and the predominant phenotype. **(F)** Triangle heatmap of cosine similarity comparing the intratumoral heterogeneity across all patients. Cosine similarity is calculated between log2 fold changes from patient-specific clusters against all other tumor cells within the respective patient. Left side annotated are patient-specific clusters from patient 1 depicted in Figure 2A and right side from patient HN01 depicted in E. We included only patients with more than 50 tumor cells.

### EMT-related transcription factor ZEB1 is highly active in metastatic epithelial differentiated OSCC cells

The transcription factors ZEB1/2, TWIST1/2, Snail (SNAI1) and Slug (SNAI2) are key for regulation of EMP [7, 27]. While mRNA expression of *SNAI2* within single OSCC cells was recently reported, the others were not detected (8). Here, we confirm this observation as we detected *SNAI2* mRNA in almost half of the OSCC cells (Figure 4A). However, detection of low expressed genes such as transcription factors by scRNAseq, especially in 10X genomics technology, becomes unreliable due to dropout effects. Also, the activity of transcription factors is often not reflected by the dynamics of their mRNA expression alone, as their activity depends additionally on protein stability and posttranslational modifications; for example, ZEB1 protein is more stable than Snail (45). To circumvent this problem, we inferred the activity of these transcription factors based on the mRNA expression profile of their target genes using the algorithm VIPER with regulons defined by DoRothEA database (30-32). Using this approach, we were able to detect high activities of ZEB1, ZEB2, Snail and Slug in OSCC cells of different patients with varying EMP phenotype (Figure 4B). Since, on the one hand, the observation that epithelial differentiation in OSCC metastases is associated with higher activity of the EMT activator ZEB1 was unexpected and, on the other hand, the method used to derive transcription factor activities appears to maybe overestimate the activity for transcriptional repressors, we addressed the plausibility of this observation (Figure 4C, Supplementary Figure 7). Consistent with the fact that one of the main functions of ZEB1 is the downregulation of E-cadherin (46) (*CDH1*), OSCC cells generally showed low expression of *CDH1* (Figure 4B). Next, we investigated the expression of ZEB1 protein by immunohistochemistry (IHC) in all 14 tumor lesions of our cohort. Since we observed nuclear ZEB1 expression in similar tumor areas as high cytoplasmic cornifin-B expression, which served as a marker of epithelial differentiation (Figure 4D, E), we validated the co-expression of ZEB-1 and cornifin-B in the same cell by immunofluorescence double-staining (Figure 4F). In line with our scRNAseq data, colocalization of both proteins was observed in a fraction of cancer cells in 9/14 (64 %) samples, double positive cells were more frequently observed in lymph node metastases (7/9, 78 %; Table 1).

**Figure 4.**
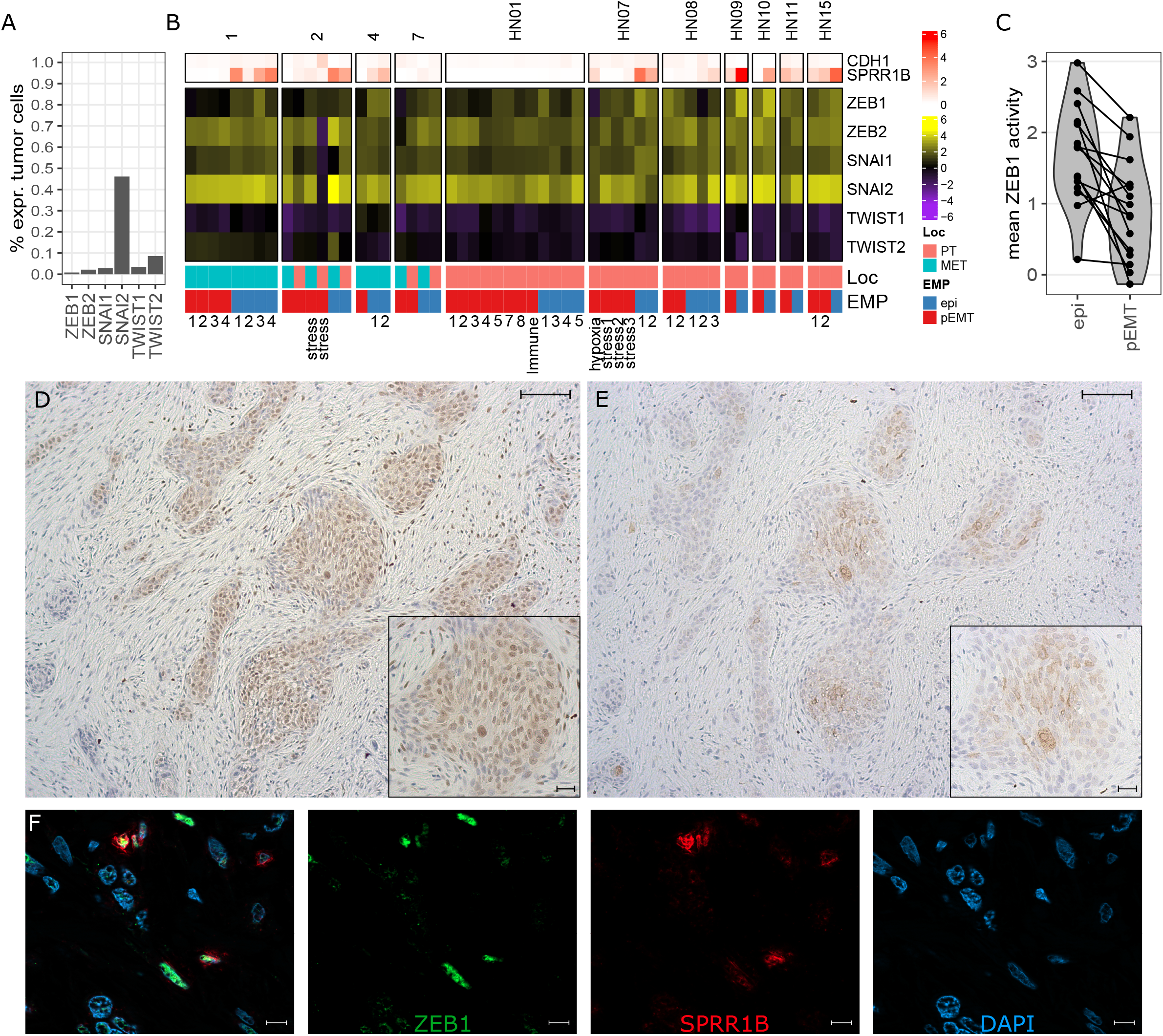
ZEB1 is highly active in metastatic epithelial differentiated OSCC cells. **(A)** Percentage of tumor cells with detectable mRNA expression from scRNAseq (more than one UMI) encoding the indicted EMP-related transcription factors. **(B)** Mean inferred activity based on the target genes of the indicated transcription factors across tumor phenotypes from EMP-related patient-specific clusters. On top the log-normalized expression of *CDH1* and cornifin-B (*SPRR1B*) is shown, on bottom the localization (primary tumor [PT] or lymph node metastasis [MET]) and respective EMP phenotype of the cluster. **(C)** Mean activity of ZEB1 for epithelial differentiated and pEMT clusters of each patient, respectively. Connecting lines show dots belonging to the same patient. **(D, E)** ZEB1 (D) and cornifin-B (E) protein expression detected in serial sections by IHC of the primary tumor from patient 2; comparable areas are depicted. Scale bars equal 200 μm in overview and 100 μm in zoomed image. **(F)** Colocalized expression of ZEB1 (green) and cornifin-B (red) detected by double staining in the lymph node metastasis of patient 1 nuclei are stained in blue (DAPI), Scale bars equal 10 μm.

## Discussion

EMT represents the reactivation of an embryonic developmental program in which cells acquire migratory and invasive properties, i.e., prerequisites for invasion and metastasis of cancer (3, 47, 48). Thus, in early stages of metastasis, tumor cells undergo EMT, whereas in established metastases the reverse process *aka* MET is also observed (49, 50). To assess the EMP-associated heterogeneity among OSCC cells and gain some insight into the dynamics of this process, we examined the transcriptomes of 7,263 individual carcinoma cells isolated from primary or metastatic OSCC. Our results not only confirm the EMP-associated heterogeneity of cancer cells in primary and metastatic OSCC, but also demonstrate progressive MET in established lymph node metastases. Interestingly, the epithelial differentiation in OSCC metastases is associated with higher activity of EMT-activator ZEB1, which was confirmed on protein level by co-expression of ZEB1 and cornifin-B in individual tumor cells by immunofluorescence staining.

EMP appears to be the main driver of cellular heterogeneity within OSCC: detailed phenotyping of cancer cells identified several clusters whose predominant functional phenotypes corresponded to different EMP states, ranging from a pEMT to an epithelial differentiated state. However, particularly the pEMT phenotypes could be overlaid by features related to angiogenesis, ECM remodeling, metabolic adaptations, stress, and interaction with the immune system. Metabolic adaptations include response to environmental limitations as hypoxia and low glucose. Low glucose conditions are counteracted with upregulation of genes related to amino acid metabolism that fuel into glycolysis (38). While previous studies suggested that the activity of specific metabolic pathways in OSCC varies widely among patients (51), we observed that hypoxia- and stress-related gene expression patterns are rather similar between patients supporting the notion of an reactive response rather than an aspect of individual tumor evolution.

In terms of EMP dynamics, it is assumed that cells may transit from one EMP state to another along a continuous spectrum of changes. Currently, however, it is also discussed if long-lived phenotypes representing discrete EMP states prevail (3, 5, 6, 8, 9, 11-13). Most studies supporting continuous transitions are based on *in vitro* or preclinical *in vivo* models that may not fully reflect the complexity of the tumor and its microenvironments (5, 6, 9). Indeed, human *in situ* or *ex vivo* studies suggested distinct EMP states; however, these approaches do not fully capture cellular dynamics (8, 12, 48). Comparing the intratumoral heterogeneity we demonstrated that within each patient, the EMP-driven heterogeneity is most prominent and epithelial differentiated phenotypes are profoundly different from pEMT clusters suggesting that these states may be rather static. Moreover, gene expression dynamics estimated by RNA velocity demonstrated epithelial differentiated cells were strongly developing towards a more pronounced epithelial differentiation with an increasing expression of genes of the epidermal differentiation complex (44). Of note, OSCC cells with a pEMT phenotype did not show such a uniform developmental direction. The assumption that epithelial differentiated metastatic cells developed later than pEMT cells, i.e., undergoing MET, is supported by an increasing number of inferred copy number gains towards increasing epithelial differentiation even if accounting for the limitations of this approach.

However, we cannot conclude whether MET happened within the metastasis or primary malignancy, as our data reflects the tumor heterogeneity within a specific timepoint of tumor evolution. Multiple disseminated tumor cells reflecting the observed EMP heterogeneity might have migrated collectively from the primary tumor that could be crucial for metastatic consolidation, as we observed a similar EMP heterogeneity also within primary tumors.

The high transcriptional activity of EMT-activator ZEB1 in epithelial differentiated OSCC cells in both primary and metastatic tumor lesions was unexpected, even considering that the scRNAseq data inferred transcription factor activities are biased towards transcriptional repressors. In the case of primary tumors this may be interpreted as the incipient EMT, but this would not work for the metastatic lesions in which ZEB1 activity was associated with a progressive epithelial differentiation. Indeed, in previous studies depletion of ZEB1 has driven tumor cells from pEMT towards an epithelial phenotype (52, 53). However, depletion of Zeb1 also reduces phenotypic variability of cancer cells particular their phenotypic/metabolic plasticity (52). While it is well established that ZEB1 together with microRNAs stabilizes EMT through a feedforward loop, this loop could also induce epithelial differentiation based on environmental factors (54). In addition to the transcriptional repressor activity, ZEB1 has been demonstrated to induce the epithelial differentiation marker cornifin-B in response to IL-1β and IFN-γ (55). We not only demonstrate the coincidence of *SPRR1B* mRNA expression and ZEB1 activity, but also the co-localization of cornifin-B and ZEB1 protein expression in OSCC lymph node metastases. Thus, although ZEB1 activity is crucial for the induction of the pEMT state, it does not seem to completely prevent partial epithelial differentiation. Remarkably, no relevant differences in *CDH1* expression were detected between the different EMP states in the metastatic OSCC lesions. Therefore, the epithelial differentiated phenotypes we observed most closely correspond to a partial epithelial differentiation analogous with the observed pEMT phenotypes. We speculate that the driving force behind this EMP-associated heterogeneity of OSCC cells is to maintain cellular integrity. For example, ZEB1 is an ATM-substrate linking ATM and CHK1, promoting homologous recombination-dependent DNA repair and thereby protecting cells from genotoxic stress whereas expression of keratin intermediate filaments helps to protect cells from stress associated apoptosis (56, 57).

## Conclusions

Single cell transcriptomics reveals that heterogeneity within OSCC cells is dominated by EMP differences resulting in distinct partial EMT and epithelial differentiated phenotypes. Particularly the partial EMT phenotypes can be accompanied by features related to metabolic adaptations, stress, and interaction with the immune system. These EMP phenotypes likely endow capabilities that are essential for the different stages of the metastatic process, including maintenance and cellular integrity. This could be a possible additional function of ZEB1, as it is also expressed during progressive epithelial differentiation in OSCC metastases.

## List of abbreviations

CNV: Copy number variation
DC: Dendritic cell
DEG: Differentially expressed genes
EC: Endothelial cell
ECM: Extracellular matrix
EMP: Epithelial-mesenchymal plasticity
EMT: Epithelial-mesenchymal transition
FFPE: Formalin-fixed, paraffin-embedded
GSEA: Gene set enrichment analysis
GSVA: Gene set variation analysis
HNSCC: Head and neck squamous cell carcinomas
HPV: Human papillomavirus
HTO: Hashtag oligo
IHC: Immunohistochemistry
MET: Mesenchymal-epithelial transition
MsigDB: Molecular signature database
OSCC: Oral cavity squamous cell carcinoma
PC: Principal component
pEMT: Partial EMT
scRNAseq: Single-cell RNA sequencing
SNN: Shared-nearest neighbor
UMAP: Uniform manifold approximation and projection

## Supplementary Information

**Supplementary Figure 1 [.pdf]:** Histology of OSCC primary and metastatic tumors.

**Supplementary Figure 2 [.pdf]:** Cell type identification by differential expression and inferred CNVs for a metachronous lymph node metastasis.

**Supplementary Figure 3 [.pdf]:** PEMT and epithelial differentiating gene expression signatures are comparable to previously published EMT signatures.

**Supplementary Figure 4 [.pdf]:** Extended analysis of the tumor phenotype characterization for the lymph node metastasis of patient 1.

**Supplementary Figure 5 [.pdf]:** Malignant phenotypes characterized across all analyzed patients.

**Supplementary Figure 6 [.pdf]:** Bulk transcriptomes reveal the cellular composition of OSCC.

**Supplementary Figure 7 [.pdf]:** Inferred transcription factor activity might be biased by activator or repressor function.

**Table 1 [.xlsx]:** Clinical and sequencing information from OSCC patients.

**Additional file 1 [.pdf]:** Supplementary Materials and Methods.

## Declarations

## Data availability

The processed datasets generated and analyzed during the current study are available in the GEO database with accession id GSE195655. The raw FASTQ files are available upon request due to privacy reasons. The high-resolution H&E Images are available through figshare under the DOI: https://doi.org/10.6084/m9.figshare.20905837.v1. The publicly available FASTQ files from Kürten *et al*. were downloaded from the sequencing read archive (SRA) accession id SRP301444.

## Code availability

Code used for analysis is available at https://github.com/sci-kai/single_cell_EMP.

## Acknowledgements

We thank Önder Bozdogan, Translational Skin Cancer Research, German Cancer Consortium (DKTK), Essen, Germany, for his histological expertise and Linda Kubat, Department of Dermatology, University Medicine Essen, Germany, for helping with microscopy. We thank Christina H. Scheel, Skin Cancer Center, Department of Dermatology, Ruhr-University Bochum, Germany, for her helpful remarks to the manuscript. We further thank Yvonne Krause and Sophia Berger, both Department of Pathology, University Medicine Essen, Germany, for their help and guidance with HTG EdgeSeq. We further thank the DKFZ Genomics and Proteomics Core Facility and the West German Genome Center in Cologne for providing Illumina sequencing and related services.

## Author Contributions

All authors approved the final version and agreed to be accountable for all aspects of the work. KH was responsible for data analysis, data curation, visualization, interpretation, conceptualization, drafting the manuscript and updating revisions. IS, LP and PG contributed by performing wet lab experiments including single-cell dissociation, scRNAseq and library preparation. FF established and performed histological, IHC and bulk transcriptome analysis. CS contributed with sample acquisition, interpretation and revising the manuscript. NS contributed by manuscript reviewing and interpretation. JG further contributed to data analysis, interpretation, and visualization. JCB was contributing by interpretation, analysis guidance, immunofluorescence examination and manuscript editing. Further, he was responsible for conceptualization and project administration including supervision and funding acquisition.

## Ethics approval and consent to participate

Written informed consent was obtained from each patient and the Ethics Committee of the Medical Faculty of the Heinrich-Heine-University Düsseldorf (#3090) approved the study. All procedures involving human participants are in accordance with the ethical standards of the institutional and research committee and with the Helsinki Declaration.

## Consent for publication

Not applicable.

