## Supplementary Figure 1 for "Mesenchymal-epithelial transition in lymph node metastases of oral squamous cell carcinoma is accompanied by ZEB1 expression"

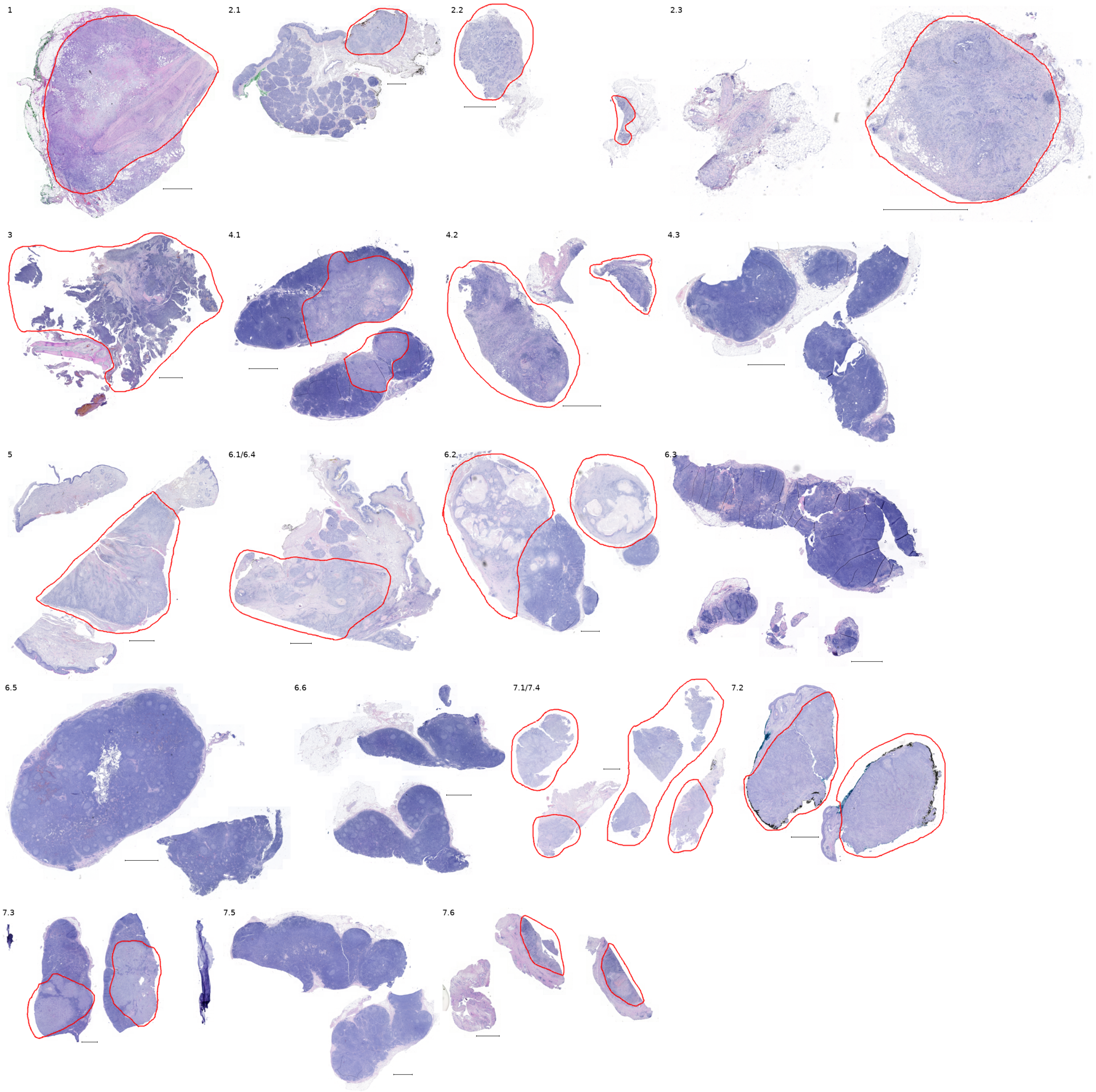


**Supplementary Figure 1 | Histology of OSCC primary and metastatic tumors.** Whole-slide image H&E staining for FFPE sections of OSCC samples. Scale bars depict 2 mm. High-resolution pictures are available through DOI: 10.6084/m9.figshare.20905837.v1.
