## Supplementary Figure 2 for "Mesenchymal-epithelial transition in lymph node metastases of oral squamous cell carcinoma is accompanied by ZEB1 expression"

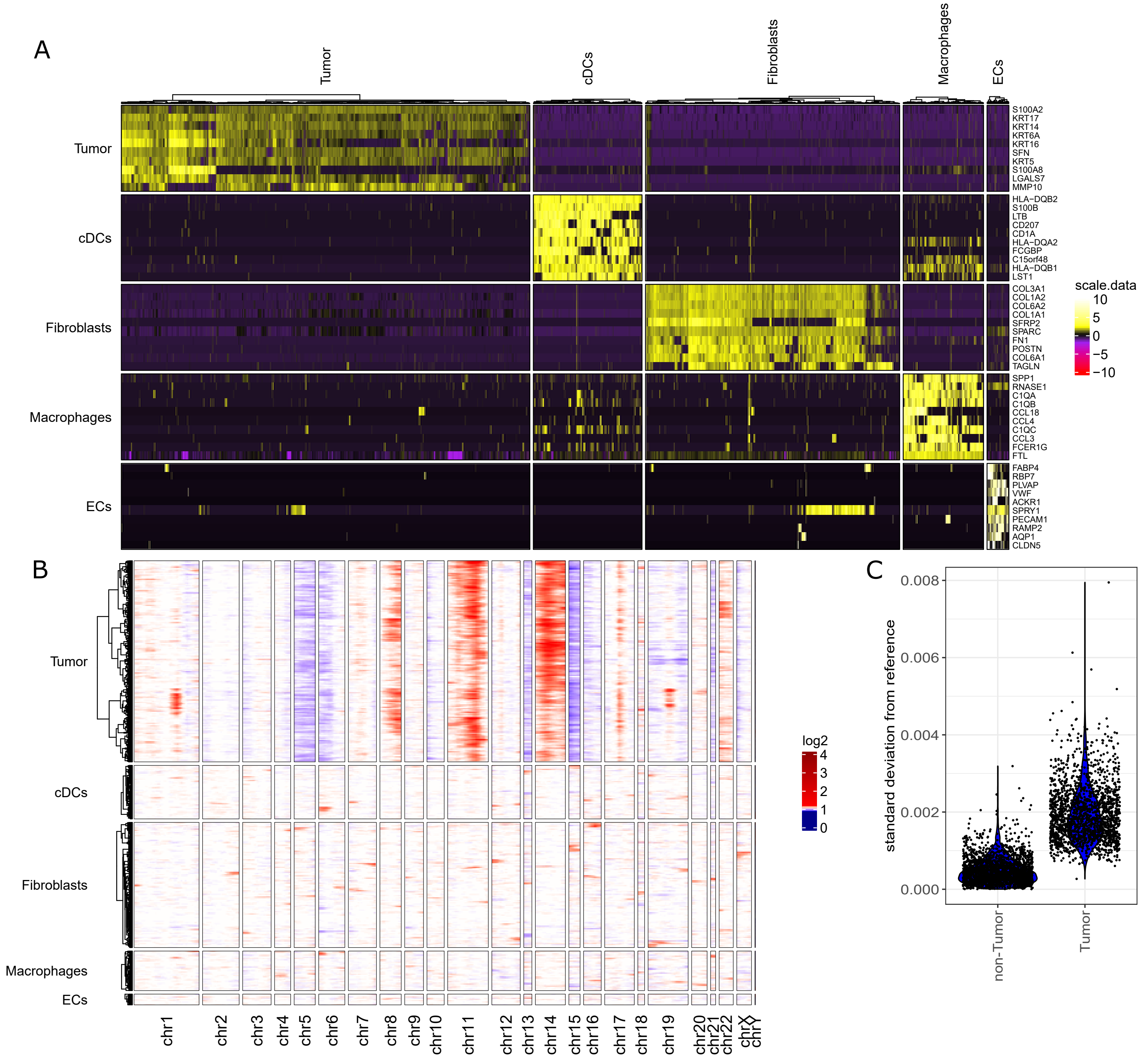


**Supplementary Figure 2 | Cell type identification by differential expression and inferred CNVs for a metachronous lymph node metastasis. (A)** Heatmap for scaled, log-normalized gene expression of all cell types from patient 1 and their top 10 DEGs (rows) against all other cells. DEGs are sorted from highest to lowest log2 foldchange. **(B)** Inferred CNVs across cells (rows) of different cell types without mitochondrial genes. Columns show genes categorized in chromosomes and ordered by genome position; hence the size of the chromosome reflects the number of detected genes and not its nucleotide length. **(C)** Standard deviation of the log2 inferCNV values to the mean of non-malignant cells compared between non-malignant and malignant cells.
