## Supplementary Figure 3 for "Mesenchymal-epithelial transition in lymph node metastases of oral squamous cell carcinoma is accompanied by ZEB1 expression"

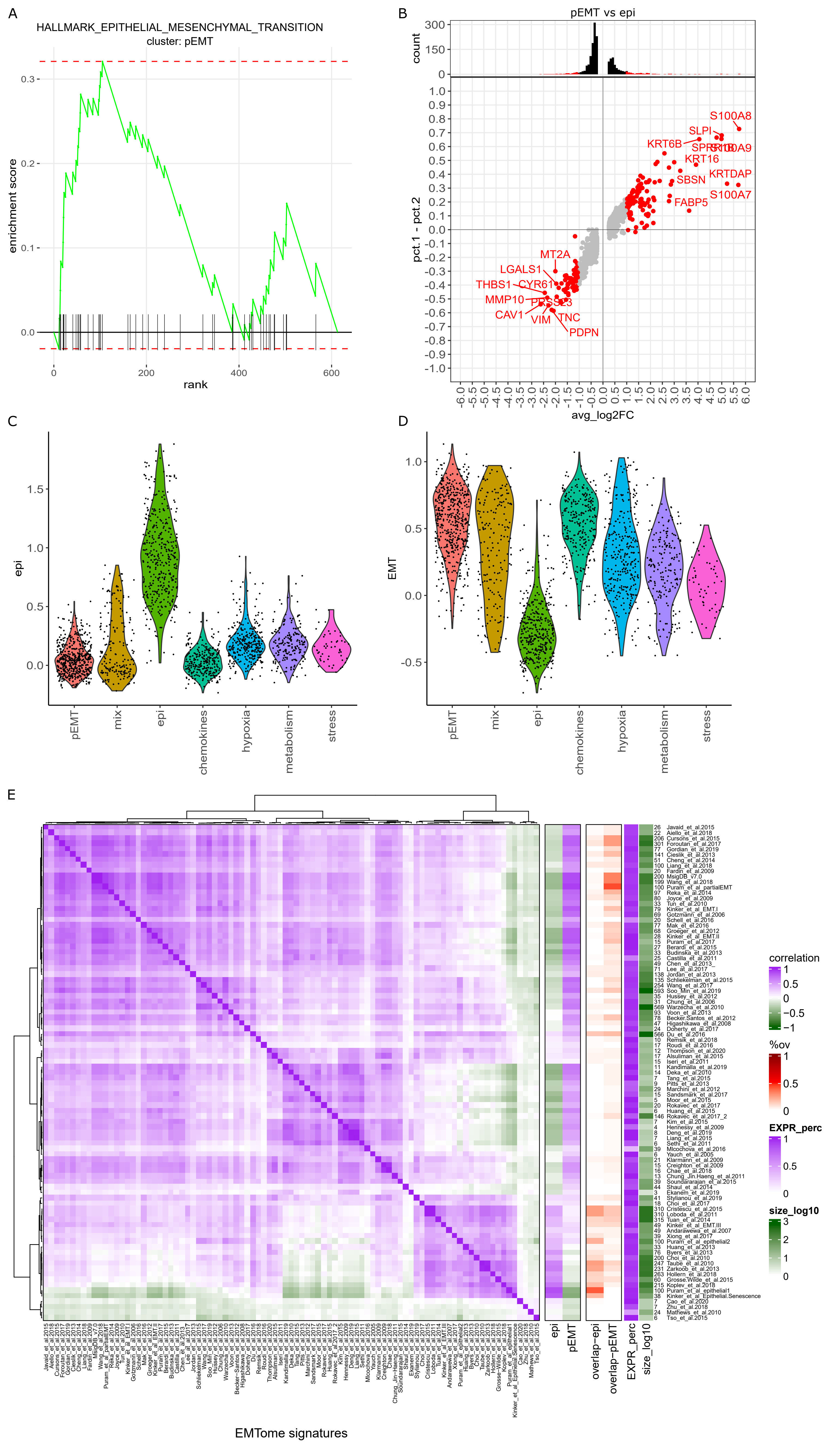


**Supplementary Figure 3 | PEMT and epithelial differentiating gene expression signatures are comparable to previously published EMT signatures. (A)** EMT hallmark gene set enrichment plot for log2 fold changes of pEMT cells against all other cells of lymph node metastasis from patient 1. Shown is the stepwise calculated enrichment score, black lines indicate genes present in the respective gene set. **(B)** Average log2 fold change of gene expression (x-axis) and differences in cellular fractions expressing the respective gene (y-axis) between pEMT and epithelial differentiated cell clusters. Labelled in red are genes with log2 foldchange below or above 1 that are included in the epithelial differentiation or pEMT signature, respectively, with top 10 genes named. The histogram on top shows the number of genes across the log2 fold change with in total 100 bins. **(C, D)** Average expression scores (y-axis) of the pEMT (C) and epithelial differentiation (D) signatures across tumor phenotypes from patient 1 depicted in figure 2A (x-axis) color-coded by these clusters. **(E)** Heatmap of correlation coefficients of GSVA scores between 91 EMP-related signatures of malignant cells, derived from the EMTome database and selected publications [7, 8, 12]. On the right side, the correlation coefficients between GSVA scores between of EMT signatures from the EMTome database and of epithelial differentiation and pEMT signatures from patient 1 (right) is shown and next to it, annotated as ”EXPR_perc”, is the fraction of genes with non-zero expression and the size of the respective EMT signature in log10 scale with the respective number next to it. Rows and columns are hierarchically clustered using a spearman correlation distance (1-cor(x,y)) and ward.D2 method.
