## Supplementary Figure 4 for "Mesenchymal-epithelial transition in lymph node metastases of oral squamous cell carcinoma is accompanied by ZEB1 expression"

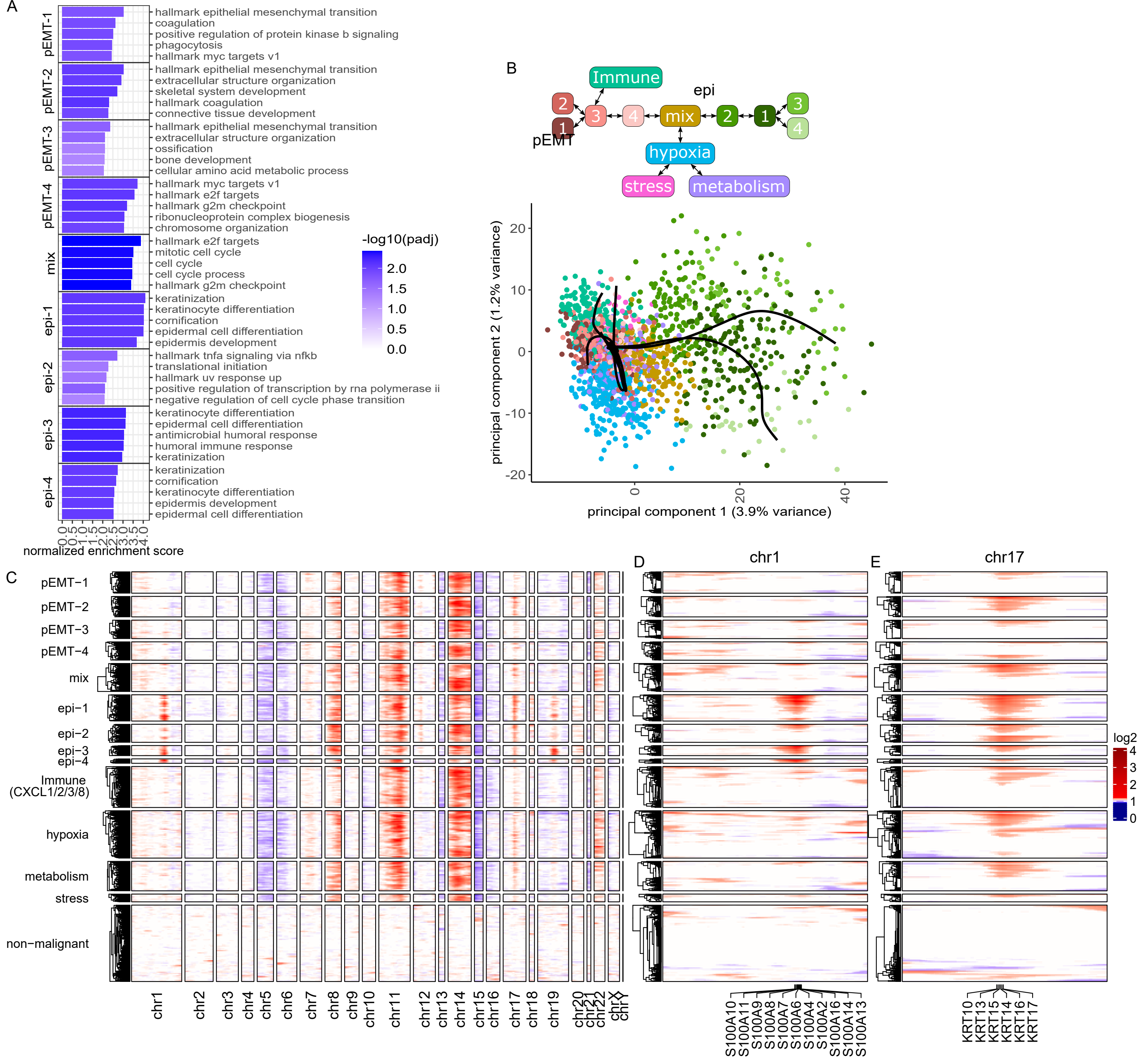


**Supplementary Figure 4 | Extended analysis of the tumor phenotype characterization for the lymph node metastasis of patient 1. (A)** Top 5 enriched gene sets from log2 foldchanges of respective tumor phenotypes by normalized enrichment scores (x-axis). Gene sets of respective phenotypes are sorted from highest to lowest enrichment. Bars are colored by the negative decadic logarithm of the Benjamini-Hochberg adjusted p-value (padj). **(B)** First two PCs of OSCC cells with all six principal curves that are derived from trajectory inference. Graph on top visualizes the relationship between malignant phenotypes, described by the principal curves forming a branching trajectory. Cells responding to environmental conditions form their own branch, indicating that the strong reactive response determines their predominant phenotype. **(C)** Inferred CNVs across tumor cells of patient 1 (rows) for all chromosomes (columns). Columns show genes categorized in chromosomes and ordered by genome position; hence the size of the chromosome reflects the number of detected genes and not its nucleotide length. Mitochondrial genes were excluded. **(D, E)** Inferred CNVs across tumor cells (rows) of chromosome 1 (D) and chromosome 17 (E) showing genes (columns) ordered by genome position. The signal on chromosome 1 is located on a genomic position on which S100 genes are accumulating and the signal on chromosome 17 on a location with accumulation of cytokeratins; most of these genes are highly expressed in the epithelial differentiated cells.
