## Supplementary Figure 5 for "Mesenchymal-epithelial transition in lymph node metastases of oral squamous cell carcinoma is accompanied by ZEB1 expression"

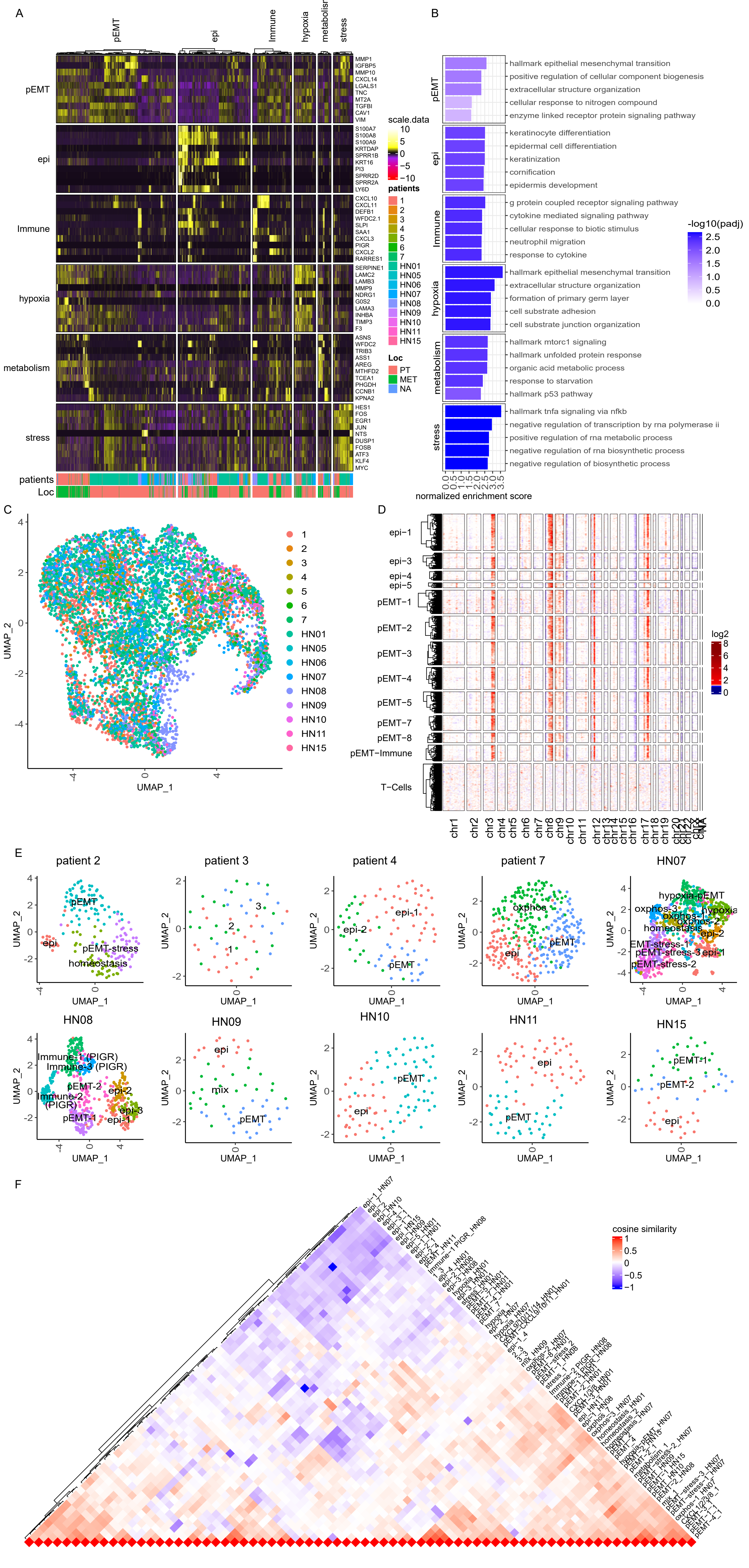

**Supplementary Figure 5 | Malignant phenotypes characterized across all analyzed patients. (A)** Heatmap for scaled, log-normalized gene expression of tumor cells (columns) split by respective phenotype depicted in Figure 3C and the top 10 DEGs (rows) of the respective phenotype against all other tumor cells. DEGs are sorted from highest to lowest log2 foldchange and row sections are ordered the same as column section. On bottom, the respective patient and localization is annotated for each cell. **(B)** Top 5 enriched gene sets from log2 foldchanges of respective tumor phenotypes by normalized enrichment scores (x-axis). Gene sets of respective phenotypes are sorted from highest to lowest enrichment. Bars are colored by the negative decadic logarithm of the Benjamini-Hochberg adjusted p-value (padj). **(C)** UMAP of OSCC cells as depicted in figure 3C with PCs corrected for patient-specific effects using harmony. Cells are annotated according to their origin. **(D)** Inferred CNVs across EMP-related OSCC cells from patient HN01 (rows) for all chromosomes (columns). Cells split by their EMP phenotype do not show any differences in their inferred CNVs pattern. Columns show genes categorized in chromosomes and ordered by genome position; hence the size of the chromosome reflects the number of detected genes and not its nucleotide length. Mitochondrial genes were excluded. **(E)** UMAPs of malignant cells from all respective patients. Cells are annotated SNN clusters and renamed according to the predominant phenotype. **(F)** Same plot as depicted in Figure 3F with the names of all patient-specific clusters as shown in E followed be the patient id.
