## Supplementary Figure 6 for "Mesenchymal-epithelial transition in lymph node metastases of oral squamous cell carcinoma is accompanied by ZEB1 expression"

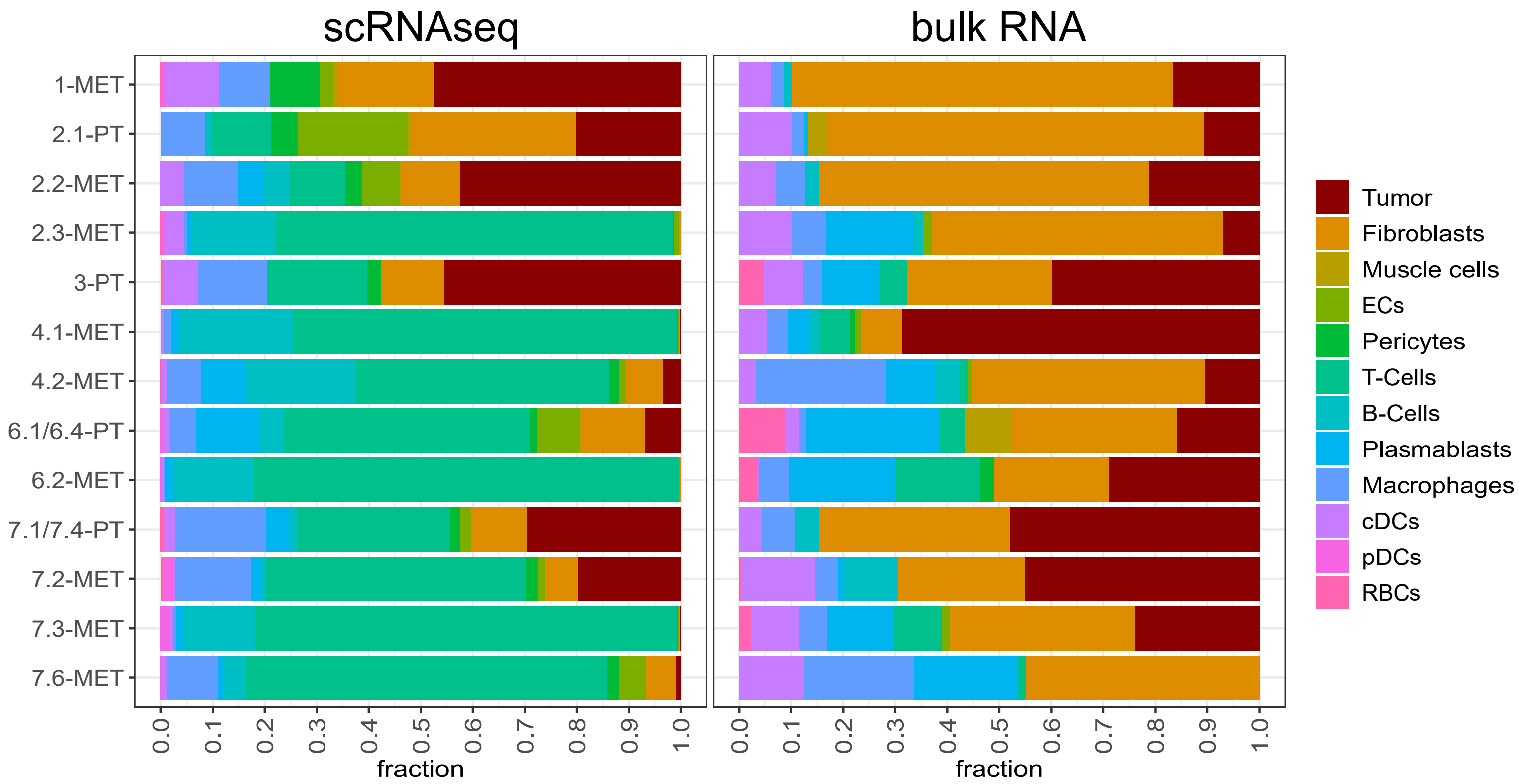


**Supplementary Figure 6 | Bulk transcriptomes reveal the cellular composition of OSCC.** Fractions of cell types across all samples including primary tumors (PT) and metastatic lymph nodes (MET) for all cells from scRNAseq (left panel) or bulk transcriptome analysis (right panel).
