## Supplementary Figure 7 for "Mesenchymal-epithelial transition in lymph node metastases of oral squamous cell carcinoma is accompanied by ZEB1 expression"

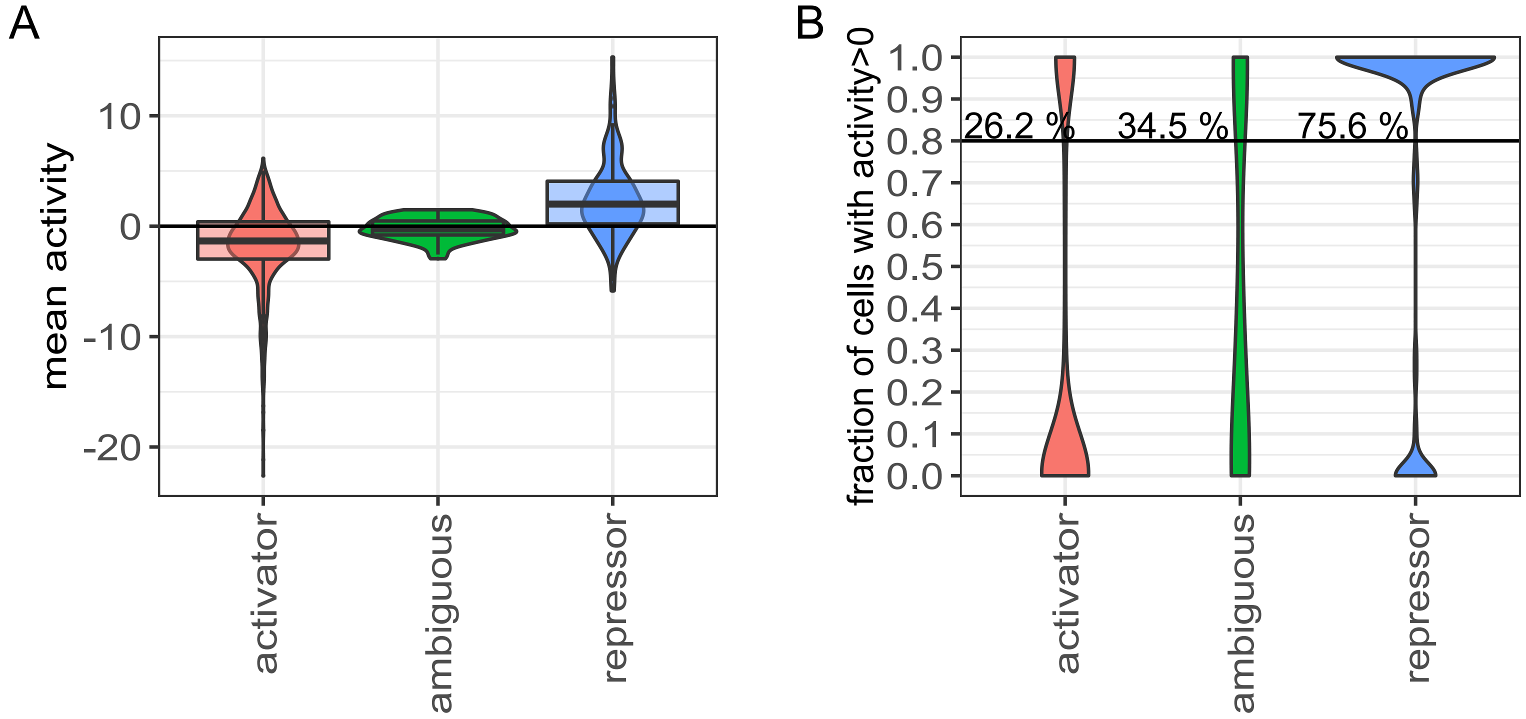


**Supplementary Figure 7 | Inferred transcription factor activity might be biased by activator or repressor function. (A)** Distribution of the mean activity of all cells from patient 1 for all transcription factors split by repressor, ambiguous and activators. Repressors and activators are defined based on more than 90% of the target genes being either repressed or upregulated, transcription factors with less than 90% for both are in the ambiguous class. **(B)** Distribution of the fraction of cells within a respective cell cluster with a transcription factor activity of greater than 0. The clusters include all cell types and malignant cell clusters from patient 1 split by activators, ambiguous and repressors. Clusters with high fraction of cells with greater than 0 activity indicate an active transcription factor, which is more prominent across repressors than for activators.
