## Additional File 1 for "Mesenchymal-epithelial transition in lymph node metastases of oral squamous cell carcinoma is accompanied by ZEB1 expression"

**Supplementary Methods**

***Cell identification***

For every sequenced library, we separately identified cells from the barcodes by evaluation of four quality criteria inspired by Luecken *et al.* [1] : (1) number of unique molecular identifiers (UMIs, nCount), (2) number of genes (nFeature), (3) percentage of mitochondrial gene expression (percent.mito) and (4) number of expressed housekeeper genes (n.exp.hkgenes), derived from Tirosh et al. without the mouse gene PRPS1L3 [2]. Accordingly, we chose manual filtering threshold for (1) and (2) s by first evaluating the histogram and combined view of both (first plot). Then, we apply the respective UMI or gene threshold and evaluate the criteria (3) and (4) by combined view (second plot). Lastly, we reevaluate each quality criteria and their combination as well as downstream analysis and adjust thresholds.

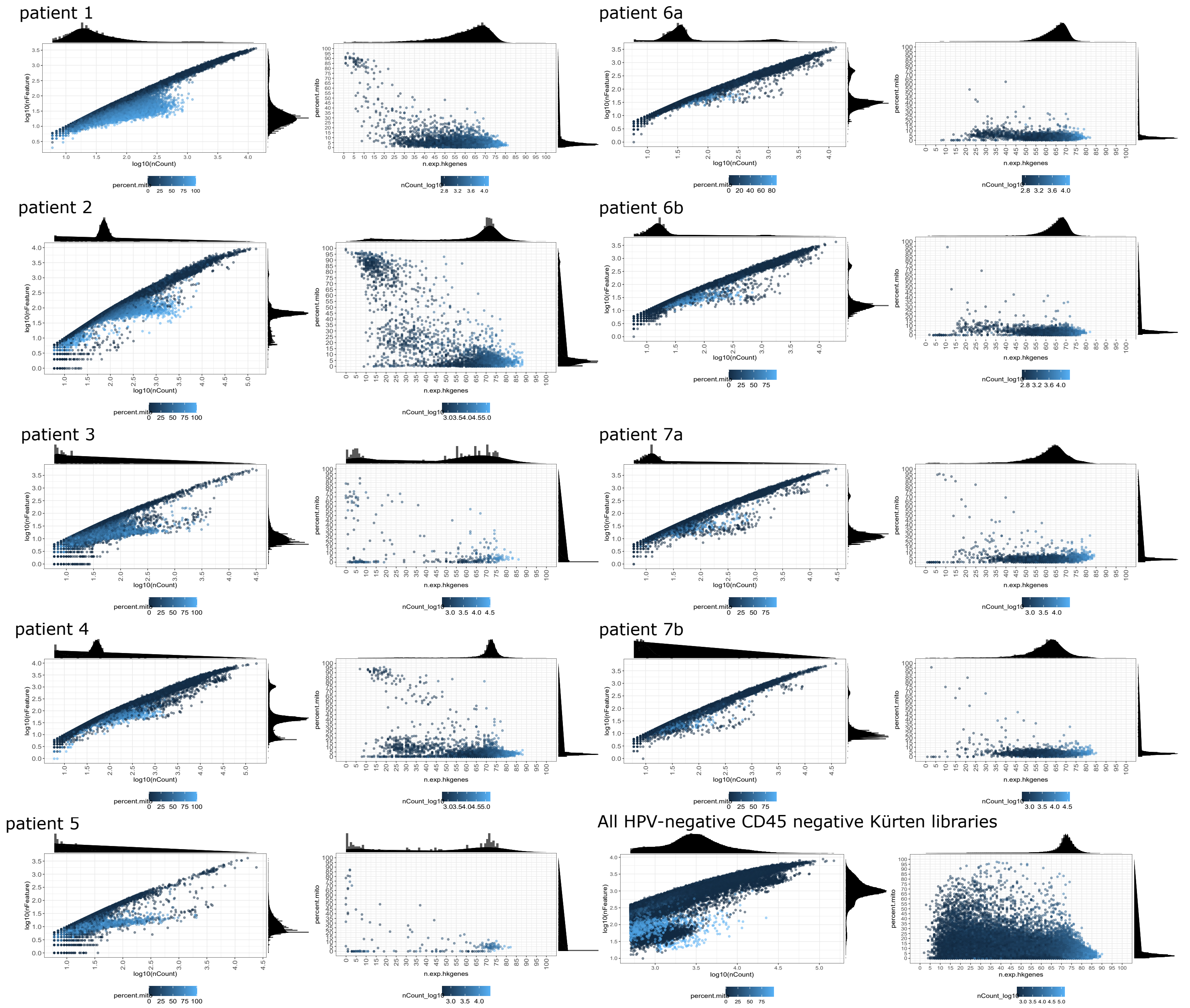


**Figure 1 | Cell identification based on quality criteria evaluation.** Quality criteria for all analyzed scRNAseq libraries. All barcodes with > 5 UMIs are shown as number of UMIs (nCount) against the number of genes (nFeature) colored by the percentage of mitochondrial gene expression with respective histograms (left plot). On the right side are all barcodes after applying the number of UMI or genes threshold showing the number of expressed housekeeper genes (maximum of 97 genes) against the percentage of mitochondrial gene expression colored by number of UMIs with respective histograms. The Kürten dataset was filtered collectively and not for every individual patient since it does not contain multiplexed hashed samples.

***Hashing***

For demultiplexing we first normalized the hashtag oligo (HTOs) expression matrix of all identified cells by centered log ratio transformation. Then, we chose manual thresholds of HTO expression for each HTO antibody and assigned samples, doublets, and cells without clear assignment. These thresholds were chosen based on combination of several criteria: We performed HTODemux from the Seurat R package once with default 99 % positive quantile for the fitted negative binomial distribution and once with the quantile greater than 50% that results the maximum numbers of singlets. The HTO expression was also visualized in scatter plots, histograms and UMAPs based on principal components to visualize the similarity of cells.

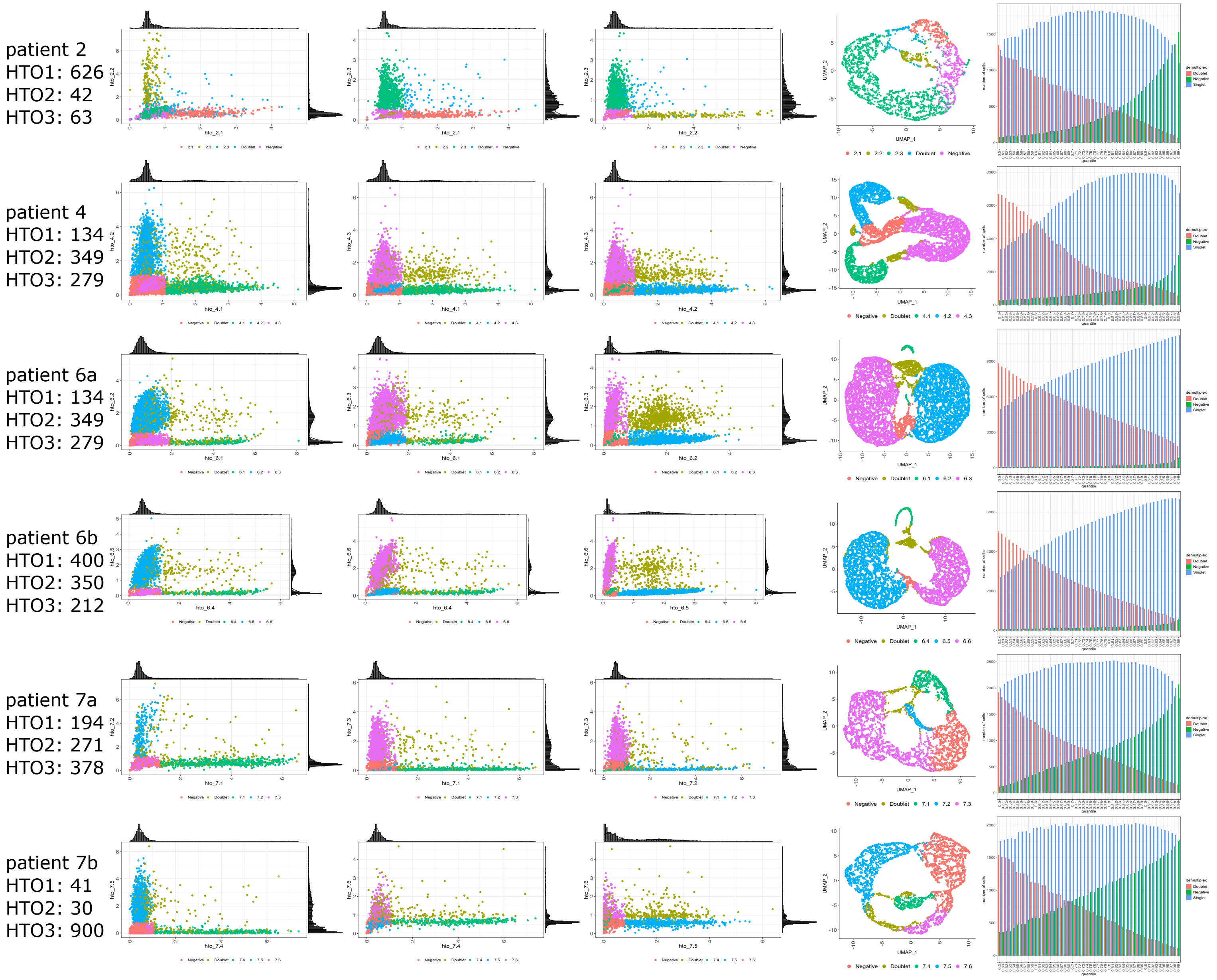


**Figure 2 | Demultiplexing of the HTO expression matrix.** The first three plots from the left show the centered log-ratio normalized expression of the three HTOs in all three different combinations. The color code shows the identified doublets, singlets and samples based on the manually determined HTO expression thresholds for demultiplexing. The fourth plot shows the UMAPs based on the principal components of the HTO expression matrix, clustering cells based on their HTO expression similarity. Color code correspond to the first three plots. The last plot on the right shows the number of cells (y-axis) identified as singlets, doublets, and negatives with different positive quantile parameter of HTODemux function from 0.5 to 0.99 (x-axis). All these information and plots were considered for choosing the manual HTO expression thresholds for demultiplexing the samples.

### *Note about trajectory and velocyto analysis:*

The trajectory derived with Slingshot yields multiple curves that reflects a branching trajectory connecting the given clusters and expressed them as multiple linear curves. Also, a single sample represents just a snapshot at a specific timepoint of tumor evolution. Therefore, the presented scRNAseq-based trajectory reflects the developmental relationships between tumor cell populations rather than ongoing evolutional processes within the sample. However, RNA velocity allows the extraction of short-term, directed dynamic information from scRNAseq data by linking the measurements to the underlying kinetics of gene expression. Hence, RNA velocity can give insights into the near future developmental processes within the sample.

### *References*

1 Luecken MD, Theis FJ. Current best practices in single-cell RNA-seq analysis: a tutorial. *Mol Syst Biol* 2019; 15: e8746; https://doi.org/10.15252/msb.20188746.

2 Tirosh I, Izar B, Prakadan SM, Wadsworth MH, Treacy D, Trombetta JJ *et al*. Dissecting the multicellular ecosystem of metastatic melanoma by single-cell RNA-seq. *Science* 2016; 352: 189; https://doi.org/10.1126/science.aad0501.
